# Precision in Spinal Cord Injury Research: A Novel Electromagnetic Impactor for a Consistent Porcine Model

**DOI:** 10.1101/2025.02.25.639133

**Authors:** Leonard Steger, Constantin Smit, Kayla Robinson, Kelley M. Kempski Leadingham, Daniel Davidar, Denis Routkevitch, Carly Weber-Levine, Kelly Jiang, Victor Quiroz, Stuart Bauer, Ruixing Liang, Ian Suk, Betty Tyler, Joshua C. Doloff, Nicholas Theodore, Amir Manbachi

## Abstract

**Purpose:** Replicating spinal cord injury (SCI) in large animals is necessary for evaluating therapeutics for potential human translation, yet there is currently no commercial, standardized device for inducing SCI. We present the fabrication and testing of a custom impactor device for producing repeatable contusion SCI in porcine models.

**Methods:** We first designed and built the device. Mechanical modeling was subsequently utilized to calibrate our benchtop testing setup. Benchtop verification was performed to measure impact force post calibration. We then used the device to generate a contusion SCI model in 2 pigs and the results were compared to an uninjured pig. Intraoperative ultrasound was used to visualize a hematoma in the injured spinal cord. Hematoxylin-eosin (H&E) and Masson’s trichrome staining were used to confirm injury presence on *ex vivo* spinal cord samples.

**Results:** Mechanical modeling forces matched benchtop impact forces within 1.4 N, indicating successful calibration of the testing setup. Our device demonstrated repeatability and the potential for modulating injury severity on the benchtop. Impactor forces were demonstrated across a range from 12.8 to 67.6 N, with variability remaining within 0.2 to 0.7 N standard deviation. The device induced two contusion injuries of different severity *in vivo*, confirmed by intraoperative ultrasound imaging and post-excision histology of the spinal cord.

**Conclusion:** Our impactor device is a major advancement towards producing repeatable and titratable contusions in large animal SCI models.

## 1 Introduction

Utilizing spinal cord injury (SCI) animal models to study the pathophysiology of SCI and evaluate potential treatments is imperative. Models of SCI can vary based on several parameters including the injury mechanism and the type of animal. The majority of human SCI results from the mechanism of transient compression, contusion, or both. [1–3]. The contusion injury mechanism in animal research is characterized by a traumatic lesion to the cord from a method such as a weight drop that causes acute injury. The compression mechanism involves a device such as a modified aneurysm clip that compresses the cord at varying force and duration [1–4]. Rodents are the most commonly studied animal model across all injury mechanisms and exhibit numerous advantages including a low cost of care, well-studied anatomy, and well-established methods for generating reliable, controlled injuries [5, 6]. One such method is the Infinite Horizons spinal cord impactor (PSI Impactors, KY, USA), which is a commercially available device for small animal SCI models [7]. Despite these benefits, successful treatments in small animal models have been shown to translate poorly to human clinical trials [5]. Among the various possible reasons for this outcome, the differences in cord anatomy and physiology between rodents and humans is a main factor. Rodents also have a propensity to recover spontaneously from SCI which makes translating outcomes to human studies and interpreting treatment effects more difficult [5, 6, 8, 9]. To overcome these issues, large animal models such as the porcine model have become increasingly studied [8, 10].

Porcine models exhibit greater translational potential due to the similarity in spinal cord anatomy and size to humans [8]. Furthermore, similarities in spinal vasculature between pigs and humans enables the study of the ischemic secondary phase of SCI. Despite these benefits, it is challenging to employ repeatable and reliably consistent methods for inducing contusion SCI models in pigs [8]. The common weight drop technique for contusion lacks consistency, as the same weight can produce different levels of injury due to issues such as friction between the weight and drop apparatus [5, 8]. Moreover, multiple impacts can occur if the weight bounces off the spinal cord [6]. Morphometric variability from measurements of spine anatomy (ex. size of intrathecal space dorsal and ventral to the cord) can also influence injury severity [11]. Additional methods such as a spring-loaded and electromagnetic impactor have been shown to enable highly controlled impacts with real time feedback of the injury parameters (Table. 1). However, there are a few limitations that demonstrate the need for continued refinement of such devices for large animal use. A reported spring-loaded device was tested with a custom operating table and must be fixed to the table, while a reported electromagnetic device was not tested *in vivo* [5, 6].

**Table 1.** Common mechanisms of generating contusion and compression type SCI in porcine models.

| Injury Type | Mechanism | Reference |
| --- | --- | --- |
| Contusion | Weight Drop | [5, 12, 13] |
| Contusion | Electromagnetic | [6] |
| Contusion | Spring-loaded | [5, 9] |
| Compression | Balloon compression | [14, 15] |
| Compression | Clip compression | [16, 17] |

Currently, there are no commercially available impactor devices to produce large animal contusion models of SCI [5]. As a result, we developed a spinal cord impactor that provides precise voltage-controlled impacts. The following work is divided into two main studies: (1) the design and development of the custom device including its specifications and fabrication; and (2) its verification and validation including calibration, benchtop verification, as well as *in vivo* ultrasound and *ex vivo* histological validation of injury. Therefore, this custom impactor is a major advancement in generating reproducible, reliable SCI in large animals.

## 2 Materials and Methods

### 2.1 Design Requirements

The general surgical workflow was considered prior to the fabrication of the device. Following a laminectomy, the device would be brought into the surgical field and the impact rod visually aligned midline over the exposed dura (Fig. 1a). The device operator would then initiate the impact, producing a contusion injury. Several design requirements were then identified based on this workflow: (1) It requires full user control and mobility of the entire system in the operating room. (2) The impact rod should be positioned in 3D space accurately for consistent placement on the dura. (3) To avoid unwanted compression and to eliminate the possibility of unwanted secondary impacts, the device should have an automatic retraction system of the impact mechanism. (4) Lastly, the device needs to be operational while maintaining surgical sterility.

**Fig. 1.**
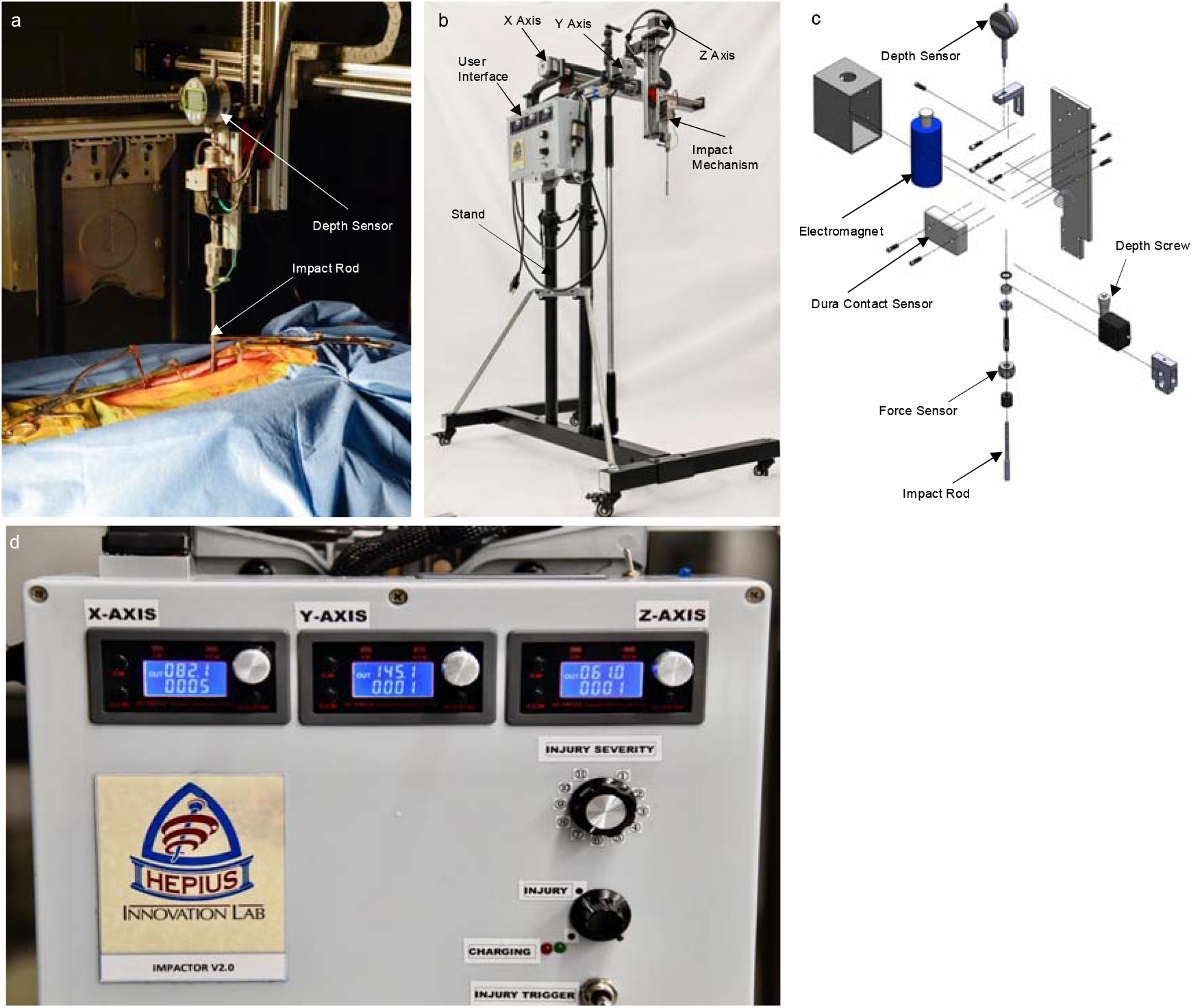
(a) Example of the impactor device in a pig model. The rod is lowered into place over the spinal cord and the depth sensor is set up to take readings. (b) Base impactor device without external depth sensor. The stand allows for portability and vertical adjustment of the device. The user interface system provides force and positional controls for the impact mechanism. (c) Exploded CAD model of the impact mechanism. The electromagnet provides the force to move the impact rod. The contact sensor alerts the user when the rod is in contact with the dura, facilitating consistent alignment above the spinal cord. (d) Controls for the user interface system. Axis knobs are positioned at the top for control over the impact rod alignment. The injury severity dial modulates impact force strength by regulating voltage through the electromagnet. Each turn of the dial (1-11) corresponds to a 10% increase in voltage through the electromagnet with a total of 12 V at 11 turns. The charging LED shines green when the device is at full power and ready for impact. The knob must be turned from the charging position to the injury position to power the impact. The injury trigger will initiate the impact.

In addition, there are several biomechanical parameters affecting injury severity that must be considered, primarily the impact force and the spinal cord displacement [5, 6, 12, 13]. To have control over both parameters, the device must be able to impact at any depth up to the full diameter of the cord and be able to impact with forces that can be modulated to inflict different levels of injury severity. Measurements on ultrasound images of the cord diameter across 2 Yorkshire pig samples revealed an average cord diameter of 5.32±0.1 mm. Previous work has shown that an impact from a 20 g weight drop causes mild injury [18].

### 2.2 Fabrication

The final impactor system (Fig. 1b-d) consists of a stand, user interface system, 3D axis arm system, and the impact mechanism. The aluminum stand can be adjusted vertically as needed and consists of a base with wheels to enable simple portability of the whole system. Fixed to the stand is a custom-built user interface box that contains axis arm controls, a dial for controlling injury severity, a charge LED indicator, and a trigger to initiate impact (Fig. 1d). Every turn of the dial (1-11 turns) changes the voltage through the electromagnet, increasing by 10% for a maximum of 12 V at 11 turns, thereby modulating impact force. Furthermore, a delay relay module (XY-WJ01, EC Technology Co., Ltd, Shenzhen, China) housed in the user interface box enables precise, automatic control over the duration of the impact. The module timer can be manually set and controls the duration of power delivered to the electromagnet as fast as in the tens of milliseconds. When the power is cut, the electromagnet plunger automatically retracts away from the spinal cord; therefore, no unwanted compression or secondary impact occurs in the event of accidental power loss.

The 3D axis arm system permits precise alignment of the impact mechanism. It is comprised of 3 linear ball screw motion modules powered by a stepper motor (FSL40, Fuyu Technology Co. Chengdu, Sichuan, China). Each arm provides controlled movement in one dimension: 500, 300, and 200 mm for the X-, Y-, and Z-axis, respectively. The impact mechanism (Fig. 1b) is attached to the Z-axis arm. The electromagnet pushes a metal rod down when the trigger on the user interface box is flipped. A depth-control screw on the side of the mechanism allows for a 0 to 7 mm range of motion of the metal rod into the spinal cord during impact. Attached to the top of the electromagnet casing is a depth sensor (C112AMXB, Mitutoyo American Corporation, USA) for real-time depth verification *in vivo* (Fig. 1c). Additionally, a contact sensor comprised of a touch switch module (TTP223, ALMOCN, USA) on the front face of the electromagnet casing will light up and set off a small alarm when the impact rod is in contact with the dura. The video demonstration in Online Resource 1 shows an example of this capability when the rod is brought in contact with a researcher’s finger. This ensures a consistent starting point of the impact. A force sensor (208C02,

PCB PIEZOTRONICS, NY, USA) is situated between the impact rod and electromagnet casing for future *in vivo* force studies. A detachable tripod stand can be mounted under the X-axis arm once all the desired adjustments have been made to stabilize the entire system and reduce variability of the impact.

### 2.3 Calibration

The goals of our benchtop verification were to assess the reproducibility of the impactor device and to ensure that it could produce similar impact forces compared to the 20 g weight drop study identified in our design requirements [18]. To address the latter, we first confirmed that our benchtop testing setup could accurately measure impact force as an evaluation of the similarity between a weight drop apparatus and our impactor device. We mechanically modeled and experimentally tested impacts with comparable parameters to the 20 g weight drop study. From this we obtained a theoretical and experimental impact force for a 20 g weight dropped at a height of 17 cm. By ensuring the theoretical and experimental values aligned, we calibrated our experimental setup and obtained a target force value for our device to generate mild injury. Ansys simulation software (R2 2023, Canonsburg, PA, USA) was used to model the benchtop testing setup with a polyurethane gel pad (AliBlue Gel Knee Crutch Pads, AliMed, Deham, MA, USA) laid on a force sensor (IMF-RSP1-020M, Loadstar Sensors, Fremont, CA, USA), using the LS-DYNA analysis system (Fig. 2a).

**Fig. 2.**
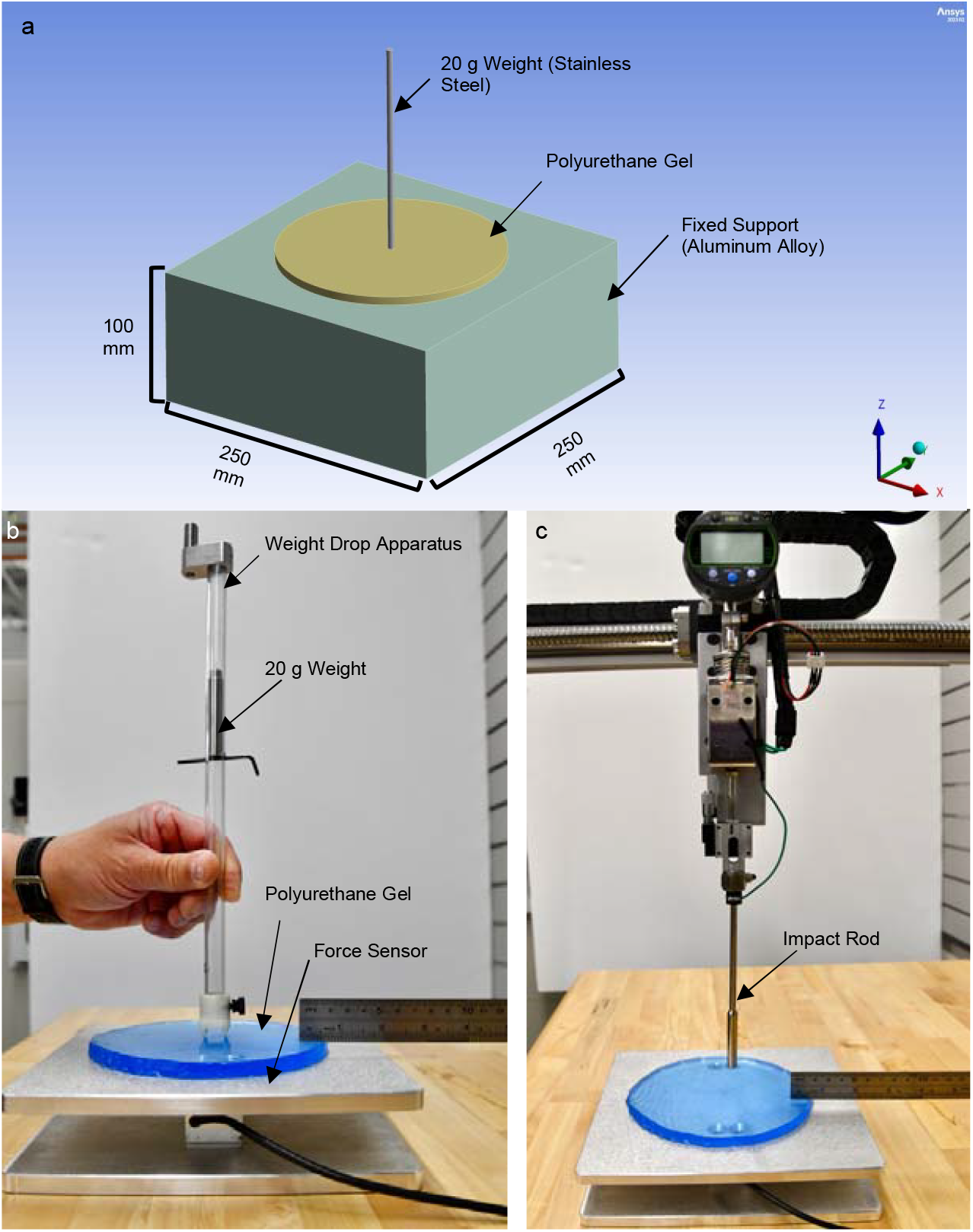
(a) Calibration setup in Ansys. A classic weight drop device was computationally modeled to provide a theoretical baseline for comparison to our device and validation of the benchtop test setup. A 20 g weight at a drop height of 17 cm was modeled due to documented comparable parameters producing a mild acute injury in porcine models [18]. A polyurethane gel was used to provide an elastic impact surface. (b) Benchtop setup for the 20 g weight drop and (c) custom impactor device with polyurethane gel placed on a metal force sensor

A 250 mm x 250 mm x 100 mm rectangular prism with a fixed top face was designated as an aluminum alloy represented our force sensor. All material mechanical properties were set to the reported value in Ansys. The aluminum alloy was set to a density of 2770 kg/m^3^ and a Young’s Modulus of 71000 MPa. The polyurethane gel cylinder was placed on top of the cube with a density of 1050 kg/m^3^ and Young’s Modulus of 6 MPa. The static friction coefficient between the gel and cube was set as 0.2, because it was assumed that the interaction between polyurethane gel and aluminum alloy would be similar to that of polyethylene and stainless-steel [19]. A steel rod representing the 20 g weight was placed on top of the center of the gel. The rod had a diameter of 5 mm and a length of 155 mm, corresponding to the dimensions of the experimental weight. The simulated drop height was the same as the experimental height at 17 cm. The mesh element size was set to 10.992 mm to ensure a balance between accuracy and computational time. The rod’s contact force with the elastic material was measured along the Z-axis, with the area of contact representing the frictional region between the table and the elastic material. Therefore, the force would be measured from the same point as in the experimental setup.

Once the modeled impact force was obtained, we dropped the 20 g weight from a handheld apparatus to get the experimental value (Fig. 2b). The drop height was 17 cm, and the peak impact force was recorded for 10 drops. The experimental setup consisted of the polyurethane-based surgical gel pad laid on our force sensor. The gel provides an elastic surface to record the weight drop impact forces. The sensor was calibrated according to the manufacturer instructions. It was then connected to a computer via a USB load cell interface (DI-1000UHS-10k, Loadstar Sensors, Fremont, CA, USA). The computer was running a high-speed single-channel load cell software (LV-1000HS-10K, Loadstar Sensors, Fremont, CA, USA) to log the force measurements.

### 2.4 Benchtop Verification

Post calibration, we characterized the reliability and strength of the impactor on the benchtop. Ten impacts per dial turn (8-11 turns) were recorded on the gel pad setup with the depth set to 3.5 mm. Settings under 8 dial turns were not included in this study since the impact force at 8 turns was already less than the 20 g weight drop test. This was repeated for depth settings of 4 mm and 5 mm. The depth setting of 3.5 mm was chosen to do an initial test with, since it would displace the cord between 50% and 100% of the cord diameter. (Fig. 2c). Depth settings of 4 and 5 mm were tested to investigate how impact force changed with depth. Impact force consistency was measured for each test, and the force values were compared to the 20 g weight drop calibration to ensure the impactor could produce a mild contusion injury. Analysis was performed to assess the statistical significance of the benchtop results. T-tests, as well as a Kruskal-Wallis test, were run to compare impact force values across the different device settings tested. The analysis was done in python using the scipy and statannotations packages [20, 21]. ChatGPT (OpenAI) was used to assist with the python code development.

### 2.5 Animal Validation

The device was evaluated in two pigs at different depth settings *in vivo* to validate its capability to produce acute contusion-type SCI and modulate injury severity. For these studies, Yorkshire swine weighing between 36 to 41 kg were used, following procedures approved by the Johns Hopkins Animal Care and Use Committee (SW23M249). Following the induction of anesthesia, the pigs were transferred to the surgical table and a posterior midline incision was performed followed by muscle dissection and a thoracic laminectomy at levels T4-T6 to expose the spinal cord (Fig. 1a).

After exposing the thoracic cord, the cavity was filled with saline and pre-injury B-mode ultrasound images were collected on a Canon Aplio i800 with a Canon i18LX5 transducer (Canon Medical Systems, Otawara, Japan). The impactor was then draped and brought into the surgical field. The impact apparatus was lowered into place using the axis arms and aligned over the exposed spinal cord at T5 (Fig. 1d). A 1.5 mg/kg rocuronium bolus was used to induce paralysis to prevent additional movement from the pig. The proper starting point on the dura was confirmed with the contact sensor. The first study tested the 10 dial turn setting at a depth of 3.5 mm to produce a mild contusion injury. The second study had depth increased to 4 mm to increase injury severity. The impact was induced by pressing the trigger on the user interface system. A short breath hold was performed for the duration of the impact to eliminate movement of the cord in relation to the impact rod. Immediately after injury the impact rod retracted automatically to prevent additional compression of the spinal cord, and the operating table was dropped in case there was any excess motion by the pig. The impactor was then removed from the surgical field, and the commercial ultrasound probe was again used to collect sagittal ultrasound images post-injury. The pre- and post-injury ultrasound images were evaluated to confirm the presence of a hematoma post-injury.

Following ultrasound imaging, the animals were euthanized at the conclusion of each study. The injured spinal cord was harvested and fixed post-excision in 4% paraformaldehyde for 24 hours. Hematoxylin-eosin (H&E) and Masson’s trichrome stains were performed on transverse slices of the tissue and imaged for analysis of the injury.

## 3 Results

### 3.1 Benchtop Calibration and Verification

The simulated and experimental impact forces from the 20 g weight drop at a height of 17 cm had a minimal difference of 1.4 N (Fig. 3a). This can be attributed to friction between the weight drop and the handheld apparatus that caused the experimental value to decrease. The close match between the experimental and simulated results verified that our experimental benchtop setup was valid, and that the force sensor was calibrated and working as intended. Furthermore, at 3.5 mm depth and 10 dial turns, the impactor displayed a similar mean impact force to the 20 g weight drop (Fig. 3a). This indicated the ability of the impactor to produce mild injury. Additionally, low variability was seen across all tests with the greatest standard deviation being 0.7 N for 9 dial turns at 5 mm depth (Fig. 3b). The maximum range for the device across all tests was between 12.8 N±0.4 N and 67.6 N±0.4 N which indicated that our device has the potential for modulating injury severity. The difference between each dial turn within a depth setting was significant so we can conclude that each dial turn represents a specific impact strength. We additionally observed an increase in impact force at the same dial turn setting with an increase in depth. However, this was primarily seen with dial turns 10 and 11, while the force at dial turns 8 and 9 was not significantly different between 4 mm and 5 mm depth. We hypothesize that this pattern showing only at less turns is a result of the stiffness of the gel pad used for benchtop testing. With lower voltage there is not enough force to penetrate as far as the depth setting allows on the gel.

**Fig. 3.**
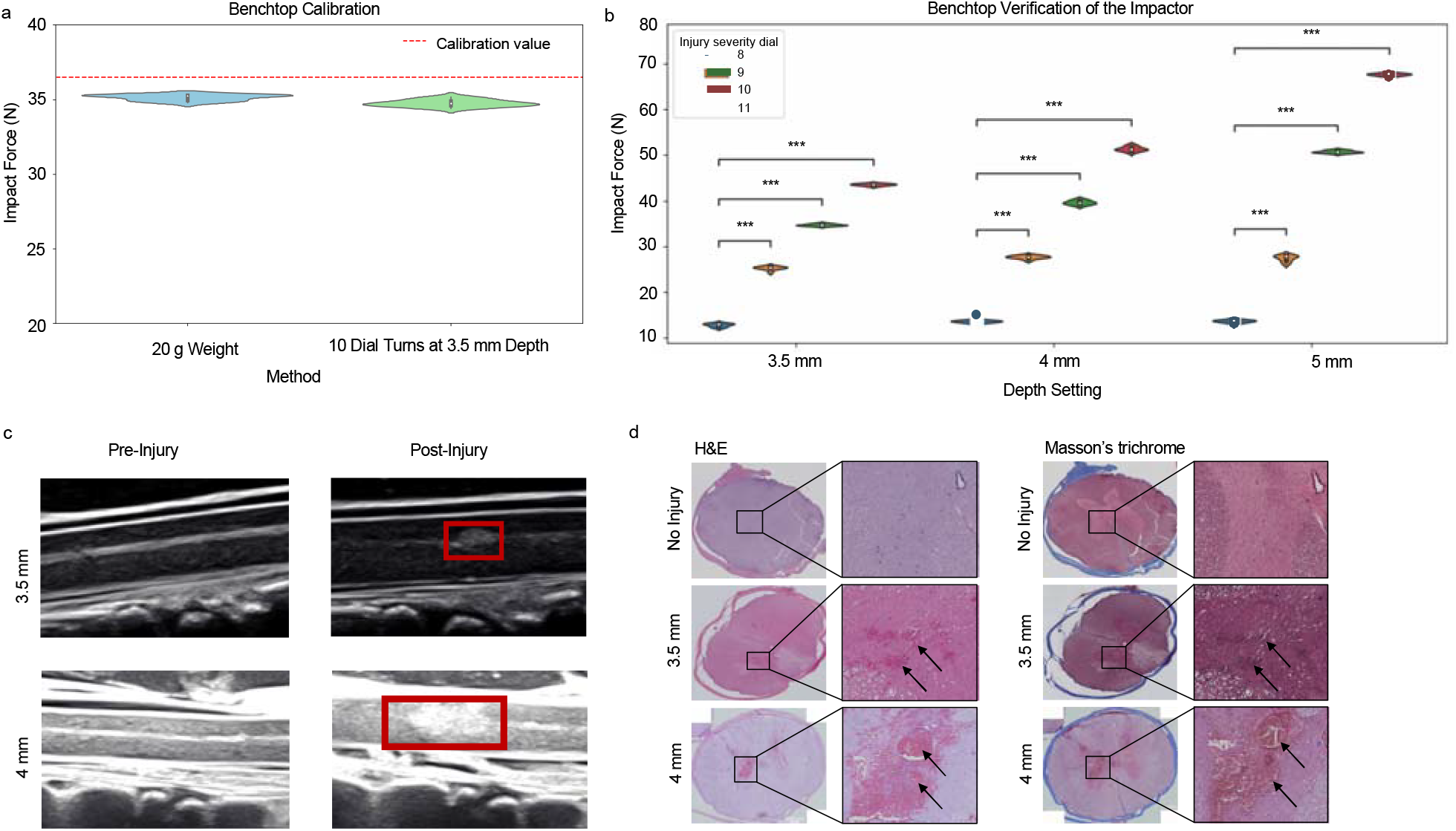
(a) Benchtop calibration revealed a mean impact force for the experimental 20 g weight drop from a height of 17 cm to be 35.1±0.2 N, while the simulated value was 36.5 N. This validated our experimental setup. Subsequent benchtop testing of the impactor device showed a mean impact force of 34.8±0.2 N at 10 dial turns and 3.5 mm depth, which indicted the device’s capability to produce mild injury. (b) Benchtop analysis for the custom impactor device. At 3.5 mm depth, the mean impact forces were 12.8±0.4 N, 25.3±0.4 N, 34.8±0.2 N, and 43.5±0.2 N for dial turns 8-11 respectively. At 4 mm depth they were 13.5±0.4 N, 27.7±0.4 N, 39.7±0.5 N, and 51.3±0.5 N for dial turns 8-11 respectively. At 5 mm depth they were 13.6±0.4 N, 27.5±0.7 N, 50.7±0.3 N, and 67.6±0.4 N for dial turns 8-11 respectively. This demonstrates the consistency of our device and the wide range of impact forces it can generate for modulating injury severity. There was a statistically significant difference between the mean impact forces for all dial turns at the same depth setting (p < 0.001). Therefore, each dial turn represents a distinct impact force. A Kruskal-Wallis test revealed a significant difference between the mean impact forces for all depth settings at the same dial turn setting (p<0.001) except between 4 mm and 5mm depth at 8 and 9 dial turns (P> 0.05). For the statistical analysis, *** represents p<0.001. (c) Pre-injury (left) and post-injury (right) sagittal B-mode images of the porcine spinal cord for the 3.5 mm and 4 mm depth setting in vivo validation studies, displaying hematoma presence post injury (red box). The injury severity dial was turned 10 times for both studies. (d) H&E and Masson’s trichrome staining were performed on the extracted spinal cord samples post-surgery for both injury tests and for an uninjured pig. At 3.5 mm depth, histological analysis of transverse slices of the cord showed injury through accumulation of red blood cells (black arrows). At 4 mm depth, there was a significant increase in the red blood cell infiltration indicated a higher injury severity. This prominent bleeding was not present in the uninjured sample

### 3.2 Animal Validation

Ultrasound imaging showed the presence of injury with visualization of a post-injury hematoma in both injury tests (Fig. 3c). The hematoma generated from the 4 mm depth test was larger compared to the injury at 3.5 mm indicating a stronger injury. H&E and Masson’s trichrome staining confirmed hematoma presence for both 4 mm and 3.5 mm tests through red blood cell (RBC) infiltration (black arrows) in the gray matter (Fig. 3d). Accumulation of these RBCs is primarily localized in the left portion of the tissue in both cases. Additionally, there was a marked increase in RBC presence in the 4 mm depth test which showed a greater injury severity. Both injury tests were compared to an uninjured tissue sample, which did not display the same cellular infiltration. This indicates our impactor can control injury severity. No gross tissue damage was observed for either test, as is ideal for the well-being of the animals.

## 4 Discussion

In this study, we addressed the need of developing a consistent, tunable impactor device for large animal models of SCI. There is a wealth of proven and available devices for small animals such as the NYU MASCIS [22] and Infinite Horizons [7] devices, yet there is no equivalent for large animal models. Towards this goal, we have developed an electromagnetic impactor device for use in pigs. We ultimately designed our device for ease of use and simple integration into the surgical workflow, while showing reproducible impact forces on the benchtop and injury severity control *in vivo*.

Key factors that were considered for ease of use were transportation, minimal assembly steps, and simple sterilization. The impactor can be transported long distances by car since the stand can be separated from the rest of the device for increased compactness. Once in the operating room, the stand can be reattached, and the device powered on. The impact rod is detachable and can be steam sterilized. The device can be draped, and the impact rod can be attached in the surgical field while maintaining sterility. We can induce a breath hold during injury, ensuring our device alignment is not affected by respiratory movement. A tripod stand can be optionally attached if increased stability is required, otherwise no assembly steps are required for use. These design choices make for an overall efficient, consistent transportation and setup pipeline prior to every surgery.

Precise control over the injury mechanism is required for repeatable impacts on the benchtop and *in vivo*. The advantage of the electromagnetic controlled impact mechanism is the ability to regulate impact duration through the relay sensor attached to the user interface box. This is necessary for replicating contusion injuries, which are characterized by a quick impact to the cord. Our device has the capability to impact with durations in the tens of milliseconds, where the impact rod automatically retracts to ensure a contusion injury without potential compression. Next to the trigger are two LED displays that signal when the electromagnet is at full charge, ensuring consistency with the power delivered to the impact mechanism. These design choices result in the minimal impact variability as seen in the benchtop analysis (Fig. 3b). Conversely, compression-type injuries are also common and are characterized by prolonged force. Our device, therefore, has the potential for inducing both compression and contusion, as the same impact duration mechanism can be extended for delayed retraction of the impact rod. Thus, our impactor device successfully replicates the contusion mechanism in a repeatable manner, with the ability for compression if desired.

Real-time feedback of the impact parameters is necessary for full characterization of the impacts produced by our device. The depth sensor allowed for confirmation of the correct depth setting on the benchtop and provides an avenue for *in vivo* readouts. Force measurements were recorded in real-time on the benchtop with an external sensor, and the internal sensor placed between the impact rod and electromagnet will be used to document impact force measurements in future animal studies. The ultrasound and histological analysis from the two injury tests (3.5 and 4 mm depth) showed modulation of injury severity, with an increase in hematoma size and RBC presence as depth increased. The injury severity dial to modulate impact force was kept at 10 turns for both tests, so spinal cord displacement was the main contributing variable for injury severity. Overall, extensive lesions and tissue damage were not present across either test, suggesting the injuries were still relatively mild. The effects of force and displacement on injury severity will require long-term survival studies that include functional outcome assessments.

We have outlined the design, fabrication, calibration, and testing of a custom large animal impactor device. Its portability and simple user interface design allow for straightforward use in large animal models. The device was able to deliver highly consistent impact forces on the benchtop and was able to tune injury severity *in vivo*. Overall, our device is a major step forward towards a reliable, and translatable option for large animal SCI model generation, that has immense potential to contribute to the large body of research on SCI.

## Supporting information

Supplemental Video 1 (Contact Sensor)

## Acknowledgements

The team thanks all Johns Hopkins Hospital veterinary staff who assisted with the animal studies.

Amir Manbachi acknowledges funding support from the National Science Foundation (NSF) STTR Phase 1 Award (#: 1938939), Defense Advanced Research Projects Agency (DARPA) Award (#: N660012024075). Denis Routkevitch acknowledges funding support from the National Institutes of Health (NIH) F30HL168823 and T32GM136577.

## Declarations

## Ethical Approval

All animal studies were completed following procedures approved by the Johns Hopkins Animal Care and Use Committee (SW23M249).

## Data availability

The benchtop study data displayed in figure 3a-b can be acquired from the corresponding author on reasonable request.

## References

[1] Young, W.: Spinal cord contusion models. Progress in brain research 137, 231–255 (2002)

[2] Sharif-Alhoseini, M., Khormali, M., Rezaei, M., Safdarian, M., Hajighadery, A., Khalatbari, M., et al.: Animal models of spinal cord injury: a systematic review. Spinal cord 55(8), 714–721 (2017)

[3] Cheriyan, T., Ryan, D., Weinreb, J., Cheriyan, J., Paul, J., Lafage, V., et al.: Spinal cord injury models: a review. Spinal cord 52(8), 588–595 (2014)

[4] Verma, R., Virdi, J.K., Singh, N., Jaggi, A.S.: Animals models of spinal cord contusion injury. The Korean journal of pain 32(1), 12–21 (2019)

[5] Züchner, M., Lervik, A., Kondratskaya, E., Bettembourg, V., Zhang, L., Haga, H.A., et al.: Development of a multimodal apparatus to generate biomechanically reproducible spinal cord injuries in large animals. Frontiers in Neurology 10, 223 (2019)

[6] Petteys, R.J., Spitz, S.M., Syed, H., Rice, R.A., Sarabia-Estrada, R., Goodwin, C.R., et al.: Design and testing of a controlled electromagnetic spinal cord impactor for use in large animal models of acute traumatic spinal cord injury. Journal of Clinical Neuroscience 43, 229–234 (2017)

[7] Scheff, S., Roberts, K.N.: Infinite horizon spinal cord contusion model. Animal models of acute neurological injuries, 423–432 (2009)

[8] Weber-Levine, C., Hersh, A.M., Jiang, K., Routkevitch, D., Tsehay, Y., Perdomo-Pantoja, A., et al.: Porcine model of spinal cord injury: a systematic review. Neurotrauma Reports 3(1), 352–368 (2022)

[9] Züchner, M., Escalona, M.J., Teige, L.H., Balafas, E., Zhang, L., Kostomitsopoulos, N., et al.: How to generate graded spinal cord injuries in swine–tools and procedures. Disease models & mechanisms 14(8), 049053 (2021)

[10] Schomberg, D.T., Miranpuri, G.S., Chopra, A., Patel, K., Meudt, J.J., Tellez, A., et al.: Translational relevance of swine models of spinal cord injury. Journal of Neurotrauma 34(3), 541–551 (2017)

[11] Kim, K.-T., Streijger, F., So, K., Manouchehri, N., Shortt, K., Okon, E.B., et al.: Differences in morphometric measures of the uninjured porcine spinal cord and dural sac predict histological and behavioral outcomes after traumatic spinal cord injury. Journal of neurotrauma 36(21), 3005–3017 (2019)

[12] Jones, C.F., Lee, J.H., Kwon, B.K., Cripton, P.A.: Development of a large-animal model to measure dynamic cerebrospinal fluid pressure during spinal cord injury. Journal of Neurosurgery: Spine 16(6), 624–635 (2012)

[13] Lee, J.H., Jones, C.F., Okon, E.B., Anderson, L., Tigchelaar, S., Kooner, P., et al.: A novel porcine model of traumatic thoracic spinal cord injury. Journal of neurotrauma 30(3), 142–159 (2013)

[14] Keller, E.E., Patras, I., Hutu, I., Roider, K., Sievert, K.-D., Aigner, L., et al.: Early sacral neuromodulation ameliorates urinary bladder function and structure in complete spinal cord injury minipigs. Neurourology and Urodynamics 39(2), 586–593 (2020)

[15] Sarwahi, V., Galina, J., Thornhill, B., Legatt, A., Pawar, A., Moguilevich, M., et al.: A study of critical events that lead to spinal cord injury and the importance of rapid reversal of surgical steps in improving neurological outcomes: a porcine model. Spine 45(4), 181–188 (2020)

[16] Zurita, M., Aguayo, C., Bonilla, C., Rodriguez, A., Vaquero, J.: Perilesional intrathecal administration of autologous bone marrow stromal cells achieves functional improvement in pigs with chronic paraplegia. Cytotherapy 15(10), 1218–1227 (2013)

[17] Kowalski, K.E., Kowalski, T., DiMarco, A.F.: Safety assessment of epidural wire electrodes for cough production in a chronic pig model of spinal cord injury. Journal of neuroscience methods 268, 98–105 (2016)

[18] Martirosyan, N.L., Kalani, M.Y.S., Bichard, W.D., Baaj, A.A., Gonzalez, L.F., Preul, M.C., et al.: Cerebrospinal fluid drainage and induced hypertension improve spinal cord perfusion after acute spinal cord injury in pigs. Neurosurgery 76(4), 461–469 (2015)

[19] Engineers Edge. https://www.engineersedge.com/coeffients of friction.htm#google vignette. Accessed: April 1, 2024

[20] Virtanen, P., Gommers, R., Oliphant, T.E., Haberland, M., Reddy, T., Cournapeau, D., et al.: SciPy 1.0 Contributors: SciPy 1.0: Fundamental Algorithms for Scientific Computing in Python. Nature Methods 17, 261–272 (2020) 10.1038/s41592-019-0686-2

[21] Charlier, F., Weber, M., Izak, D., Harkin, E., Magnus, M., Lalli, J., (2022). Statannotations (v0.6). Zenodo. 10.5281/zenodo.7213391

[22] Gruner, J.A.: A monitored contusion model of spinal cord injury in the rat. Journal of neurotrauma 9(2), 123–128 (1992)

